# Response of whole plant water use to limiting light and water conditions are independent of each other in seedlings of seasonally dry tropical forests

**DOI:** 10.1101/2020.09.16.299297

**Authors:** Ron Sunny, Anirban Guha, Asmi Jezeera, Kavya Mohan N, Neha Mohan Babu, Deepak Barua

## Abstract

How co-occurring species vary in the utilization of a shared and limited supply of water, especially in the context of other limiting resources like light, is essential for understanding processes that facilitate species coexistence and community assembly. For seedlings in a seasonally dry tropical forest that experience large heterogeny in light and water conditions, how water use, leaf physiology, and subsequently plant growth, is affected by limited water and light availability is still not well understood. In a controlled common garden experiment with four co-existing and commonly occurring dry tropical forest species, we examined how whole plant water uptake, responds to limiting water and light conditions and whether these responses are reflected in leaf physiology, and translated to growth. Water use varied dramatically in seedlings of the four species with a five-fold difference in well-watered plants grown in full sunlight. Species varied in their response to shade, but did not differ in responses to the low water treatment, possibly resulting from the strong selective force imposed by the very low water availability and the long dry period characteristic of these seasonally dry forests. Interestingly, species responses in water use, physiology, and growth in limiting water conditions were independent of light. Thus, species response to both these limiting conditions may evolve independently of each other. Responses in water use were largely congruent with responses in leaf physiology and growth. However, while magnitude of changes in leaf physiology were largely driven by light conditions, changes in whole plant water use and growth were influenced to a greater degree by the water treatment. This highlights the need to measure whole plant water use to better understand plant growth responses in these seasonally dry tropical forests.

## Introduction

Growth, reproduction and survival of plants strongly depends on the ability to acquire and utilize water (Santiago et al. 2004, Poorter et al. 2010, Fan et al. 2012). There is large variation in species water use (Wullschleger et al. 1998, Ehleringer et al. 1999, Collins and Bras 2007) and quantifying this variation, understanding how it relates to leaf level physiology, and translates to growth is important to understand competition between co-occurring species, coexistence and community assembly. Additionally, for tropical forests seedlings that have to cope with heterogeneous forest structures and light environments, in addition to seasonal rainfall regimes, it is important to understand how water use responds to varying water and light conditions (Kitajma et al. 1994, Holmgen et al, 2012). This becomes particularly important given current changing rainfall patterns and increased fragmentation due to climate change and anthropogenic disturbances (Feng et al. 2013, IPCC 2014).

Light is an important limiting factor that is often negatively correlated with water availability in many systems including tropical forests. Large statured trees with dense canopies in wet forests result in diminished light availability in the understory, and conversely, low plant cover and open canopies allow greater availability of light in dry forests (Kitajma et al. 1994, Johnson et al. 2017, Comita et al. 2014). It has been proposed that plants face a trade-off in responding to limiting water and light as a result of contrasting physiological and morphological adaptations required for dealing with each of these limiting resources separately (Smith and Huston 1989, Holmgren et al. 2012, Kupers et al. 2019). However, results from other studies suggest that plant responses to limiting water may be independent of light (Sack et al. 2004, Sánchez-Gómez et al. 2006). In fact, morphological and physiological adaptations can simultaneously reduce requirements for both light and water, and this would allow greater drought and shade tolerance (Holmgren 2000, Holmgren et al. 2012), and coordinated whole plant resource acquisition (Reich 2014). Thus, growth responses to limiting water and light availability are still not understood and could be negatively related (trade-off between responses to limiting water and shade tolerance), positively related (facilitation and coordinated resource acquisition), or independent of each other.

The primary mechanisms that might explain a trade-off between resistance to low water and tolerance to shade is with respect to differential allocation to leaf and root. An increased leaf area fractions and a corresponding decrease in root allocation would increase the efficiency of light capture but might also increase sensitivity to low water availability (Smith and Huston 1989). In contrast, conservative strategies might select for traits which result in a reduced demand for resources including light and water (Reich 2014). This might allow plants to overcome potential trade-offs to exhibit coordinated whole plant resource acquisitions strategies and positive interspecies relationships between responses to limiting light and water. This understanding of the relationship between responses to limiting water and light has important implications for niche partitioning and species interactions in habitats with spatial and temporal variation in light and water availability.

Water use by plants involves uptake by roots from the soil, flux through the stem and leaf xylem vessels and transpiration loss through the stomata to the atmosphere. Studies of water use that have quantified stem xylem conductance (He et al. 2020), leaf xylem conductance (Nardini and Luglio 2014), and leaf stomatal conductance (Cavaleri and Sack 2010) suggest that water use is important in understanding association with habitat specific water availability. However, inferences from these studies may be limited because quantification of water use is confined to scales that are smaller than the whole plant level (Meinzer et al. 2010). Organ-level studies may not be able to account for the ability to mitigate limiting water availability by other means including deep roots (Paz 2003, Pérez-Ramos et al. 2013, Paz et al. 2015, Brum et al. 2019), minimizing total leaf area (Bucci et al. 2005), or using stem water reserves (Wolfe and Goldstein 2017). A more comprehensive understanding of how water use responds to limiting resource availability requires whole plant level integrated studies (Reich 2014).

Light and water are key abiotic factors limiting growth and establishment of seedlings in seasonally dry tropical forests like our study system in the Northern Western Ghats range of peninsular India (Pascal 1988). Water availability in this region is highly seasonal with a long dry period that extends for eight months (CRU TS version 4.1). There is large spatial heterogeneity in forest cover which ranges from open savanna like matrices with less than 30% cover, to closed canopy tall statured forests with 100% cover. Thus, these systems have spatially and temporarily variable light and water conditions that may allow co-existence of trees with differing water use strategies (Sterck et al. 2011). In this study, we examined variation in whole plant water uptake in experimentally grown seedlings using the gravimetric method to overcome limitations of organ level proxies. For seedling of four commonly occurring and co-existing dry tropical forest tree species, we examined: 1) How water uptake responds to limiting water and light conditions; 2) If species responses in water uptake to the light and water treatments are reflected in leaf physiology and whole plant biomass allocation; and, 3) If these responses are translated to growth.

## Methods

### Species selection and study area

The study species were selected from a set of 80 species commonly found in seasonally dry tropical forest of the Northern Western Ghats range, India (19.1320°N, 73.5540°E). Based on data for leaf functional traits, and leaf phenology (Barua unpublished), we identified representative species that span the range of leafing behaviour and leaf functional trait values observed in this region, and selected the following species for this experimental study: *Terminalia chebula* and *Diospyros melanosylon* are relatively fast growing deciduous species with low leaf mass per area (LMA) and high leaf nitrogen content (LNC), *Syzygium cumminii* is a semi-evergreen species with moderate LMA and LNC, while *Memecylon umbellatum* is a slow growing evergreen species with high LMA and low LNC.

The climate in this region is highly seasonal, and most of the annual rainfall of 2266mm falls between June and October. The long dry period extends from November to May. The landscape is topographically diverse with valleys carved out by rivers and their tributaries. The top of these valleys consist of flattened ridges. Soil depth varies from very shallow (< 10cm) in these flattened ridges to relatively deep (>100cm) in the mid slopes and valleys.

These ridge tops (henceforth open habitats) with low soil depth, low water availability and high light availability are characterized by scrub/savanna vegetation consisting of stunted trees, shrubs and lianas with a high percentage of deciduous species. At the other extreme the valley forests (henceforth closed habitats) with greater soil depth, higher soil water availability, and low light availability are dominated by tall statured evergreen tree species. The transition zones on the slopes (henceforth edge habitats) consist of the smaller patches of fragmented forests that contain a mixture of evergreen and deciduous species with some overlap with both the open and the closed forests. Tree heights in the open habitats range from 3-5 m, in the edge habitats from 10-15 m, and in the closed habitats from 15-20 m.

### Common garden experimental design and light and water treatments

The common garden experiment was carried out in an experimental plot at the Indian Institute of Science Education and Research, Pune, India. Seedlings of the four study species were obtained from a local nursery, J.E. Farms in February 2015. These 2.5 year old seedlings were transferred to 19 Litre mini-lysimeter pots (20 cm diameter and 60 cm length) with 18 kg of uniformly mixed dry red alfisol (pH 7.2) supplemented with organic manure (1:50 v/v) and urea (0.05 g·kg^−1^ soil). Seedlings were allowed to acclimate to the lysimeter pots for a period of 30 days. A full factorial design of two levels of light (open sun and shade), and water (control and low water) was imposed for a period of 50 days. We selected 15 % of full sunlight, and 40 % of saturated pot water content as the limiting light and water treatments, respectively. These light and water levels were chosen to impose limiting but non-lethal levels based on a literature survey of previous experiments and also to simulate light in shaded habits and soil moisture content in the early dry season in our study site. Six replicate individuals of each species were used for each species per treatment giving 24 individuals per species and a total of 96 individuals in the experiment.

The shade treatment was imposed using a nylon net that allowed approximately 15 % of the sunlight to pass through, thus replicating the light conditions observed in shaded habitats in the field. Plants subjected to the open sun treatment were kept in the open in full sunlight. The average photosynthetic photon flux density at midday in the shade treatment was 153.7 ± 68 μmol·m^-2^·sec^-1^, and in the open sun treatment was 1109.3 ± 440 μmol·m^-2^·sec^-1^. For the watering treatments, control pots were maintained at 80 % of the saturated pot water content, while low water treatment pots were maintained at 40 % of the saturated pot water content.

Saturated pot water content was estimated for a subset of the pots at the beginning of the experiment. Pots were weighed at regular intervals (usually every 3 days) during the experiment to determine the amount of water lost in that period and to bring the pot water back to the appropriate treatment levels. A thin plastic sheet was taped to the rim of the pot and tied to the base of the stem of the seedlings to prevent water loss from the soil. Soil moisture content (SMC) was quantified by collecting a soil core from the small holes made on the sides of the pot. Soil cores were weighed for fresh weight, oven dried for three days at 95 °C, and the dry weighed measured. SMC was calculated as: SMC= (FW-DW)/DW.

### Quantification of plant water uptake, leaf physiology, biomass allocation and growth

Whole plant water uptake was quantified by weighing all the individual pots every three days during the experiment. Leaf gas exchange parameters were quantified with a LICOR, LI-6400XT Portable Photosynthesis System (LI-COR, Lincoln, USA) using the standard broad-leaf cuvette (6 cm^2^) fitted with the LICOR-6400-02B LED light source on the first fully expanded and mature leaf of every individual between days 44 to 49 after imposition of the treatments. All gas exchange measurements were made between 9:00-11:00 hrs with the cuvette light, chamber CO_2_ concentrations (incoming reference), relative humidity and temperature set at 1400 μmol·m^2^·s^-1^ PPFD, 390 + 10 ppm, 50-60 %, and 28-30 °C, respectively. Stem height and stem diameter were measured for all individuals at the start and at the end of the treatment period to quantify growth. Height was measured with a measuring tape from the base of the stem to the tip of the terminal branch, and stem diameter was measured using a vernier caliper at a height of 10cm from the base of the seedling. All plants were harvested at the end of the treatment period to quantify biomass fractions of root, leaf and stem. Leaf area was estimated for all individuals. Leaves that had emerged during the experiment period were marked and this allowed us to measure leaf area increase during the treatment period. Leaf, stem and root tissue were oven dried at 70°C for 3-5 days to constant weight to quantify dry mass.

### Statistical analysis

All responses measured were tested for normality and transformed when necessary. The response of the four study species to the light and water treatments were analyzed using a full factorial ANOVA with species, light and water as fixed effects. All analysis were performed with Statistica (version 9.1, Statsoft, Tulsa, OK, USA).

## Results

Whole plant water uptake varied nearly five-fold among seedlings of these four co-existing species. Water uptake in the well watered control treatments in full sunlight ranged from around 200 ml per day for *Memecylon* to nearly 1000 ml per day for *Syzygium* (Fig. 1). In addition to inherent differences in water uptake, the species responded differently to the light and water treatments over the course of the experiment.

**Figure 1:**
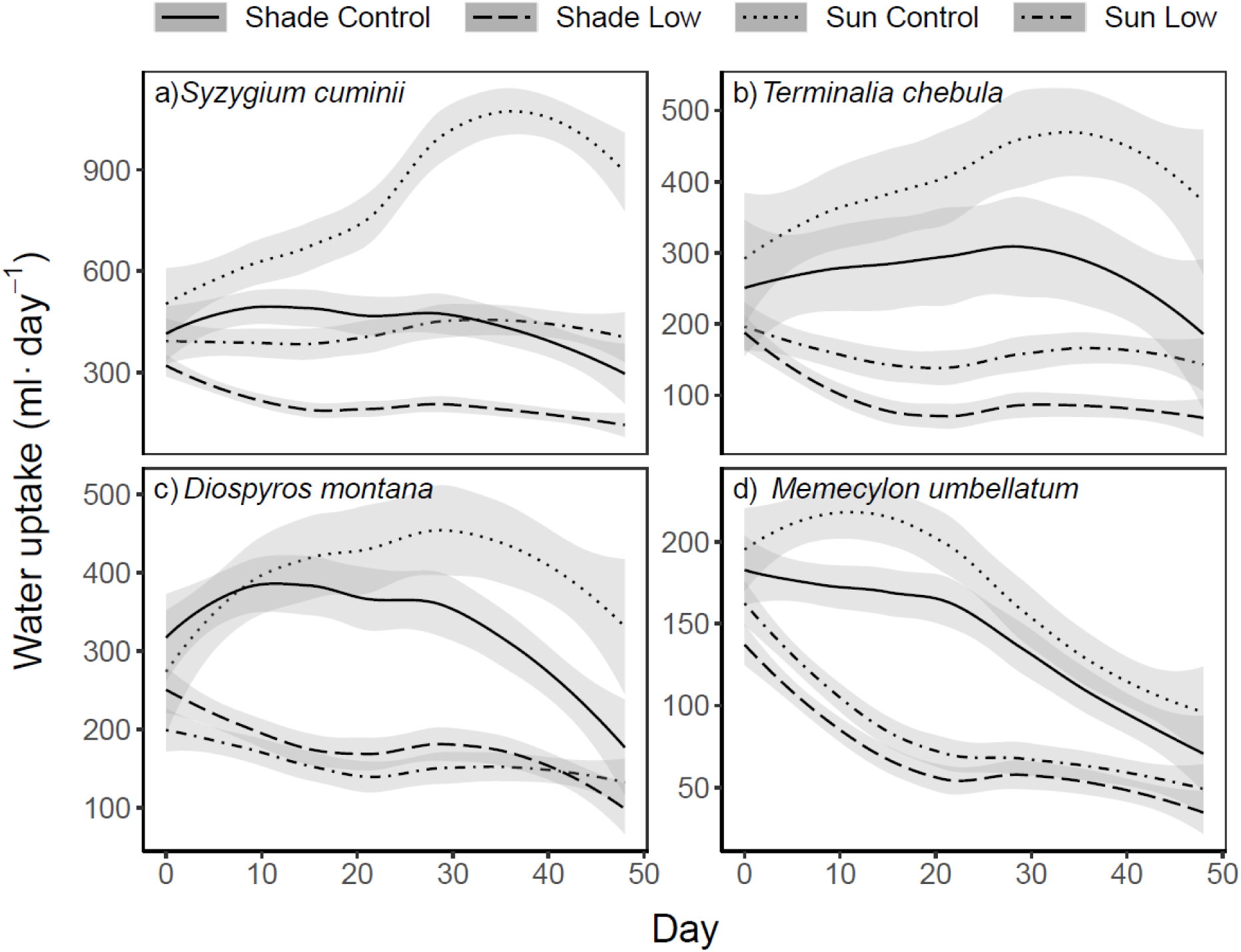
Whole plant water uptake over the course of the experiment in response to light and water treatments in seedlings of the four study species: a) *Syzygium cumini*; b) *Terminalia chebula*; c) *Diospyros montana*; and, d) *Memycelon umbellatum*. The lines represent a local polynomial regression fit (loess) to the whole plant water uptake per day quantified for six individuals of every species and treatment combination. The grey bands around the lines represent 95% confidence intervals. The light and water treatments are depicted by: shade control - solid lines; shade and low water - dashed lines; sun control - dotted lines; sun and low water-lines with alternating dots and dashes.

Water uptake for six individuals for each species and treatment was quantified for fifty days. As expected reduced water availability resulted in a decrease in whole plant water uptake and this was consistent across all four species (Fig. 2a-d, Table 2). Again as expected, water uptake was lower for plants in the shade than in the open sun. The reduction in water uptake in the low water treatment was similar for both sun and shade plants indicating that the shade treatments did not alleviate effects of reduced water availability (Table 2). The reduction in water uptake between the open sun and shade treatments differed among species (Table 2), and was lowest for Memecylon and highest for Syzygium.

**Table 1:**
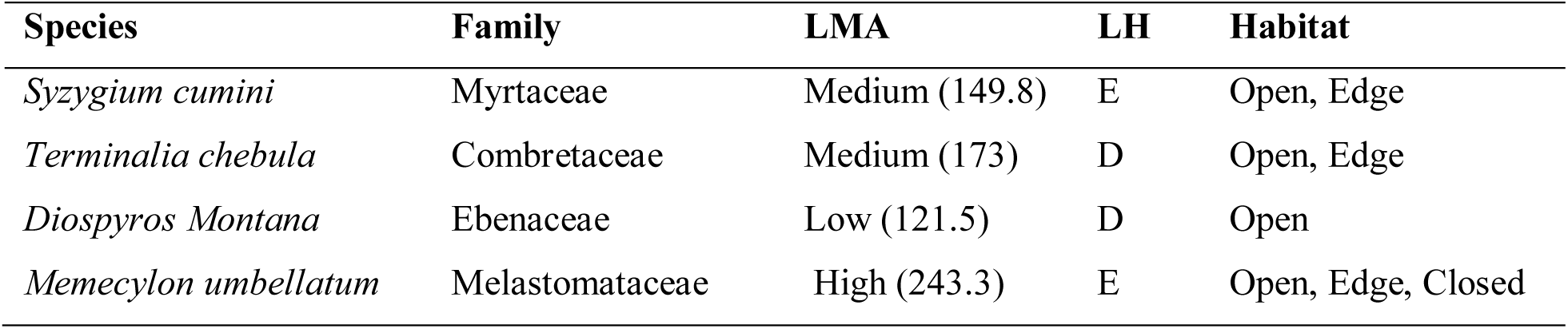
The study species with taxonomic affiliation, leaf mass per area (LMA, g·m^-2^), leaf habit (LH; E - evergreen, D - deciduous) and preferred habitat in seasonally dry tropical forest in Nigdale, Maharashtra in the Northern Western Ghats of India. LMA was classified as low, medium and high based on the distribution of traits values across 80 species in these forests.

| Species | Family | LMA | LH | Habitat |
| --- | --- | --- | --- | --- |
| *Syzygium cumini* | Myrtaceae | Medium (149.8) | E | Open, Edge |
| *Terminalia chebula* | Combretaceae | Medium (173) | D | Open, Edge |
| *Diospyros Montana* | Ebenaceae | Low (121.5) | D | Open |
| *Memecylon umbellatum* | Melastomataceae | High (243.3) | E | Open, Edge, Closed |

**Table 2:** Results for 3-way ANOVA for the effects of light treatment (L), watering regime (W), species (S) and their interactions for: a) water uptake; b) physiology; c) biomass allocation; and, growth. F statistic are presented and statistical significance indicated in bold and by the presence of asterisk (* *P* < 0.05, ** *P* < 0.01).

| Processes | Light | Water | Species | LxW | LxS | WxS | LxWxS |
| --- | --- | --- | --- | --- | --- | --- | --- |
| **a) Water Uptake** |  |  |  |  |  |  |  |
| Per individual | **37.2\*\*** | **170.1\*\*** | **85.9\*\*** | 0.0 | **6.2\*\*** | 0.9 | 0.8 |
| Leaf area normalized | **79.8\*\*** | **86.9\*\*** | **15.5\*\*** | 3.6 | **3.9\*** | 1.9 | 0.1 |
| **b) Physiology** |  |  |  |  |  |  |  |
| Photosynthesis | **62.6\*\*** | **4.7 \*** | **21.8\*\*** | 1.5 | **9.4\*\*** | 0.4 | 1.1 |
| Conductance | **18.0\*\*** | **5.8 \*** | **12.6\*\*** | 0.2 | 2.1 | 0.0 | 0.4 |
| Transpiration | **8.5 \*** | **10.3 \*** | **14.2\*\*** | 1.7 | **2.6\*** | 0.4 | 0.2 |
| **c) Biomass allocation** |  |  |  |  |  |  |  |
| Leaf mass fraction | 3.2 | **11.2\*\*** | **20.3\*\*** | 0.7 | 0.6 | **3.7\*** | 1.3 |
| Stem mass fraction | 0.9 | **4.3 \*** | **6.5\*\*** | 0.0 | 1.1 | 0.7 | 0.9 |
| Root mass fraction | 0.6 | **38.9\*\*** | **27.8\*\*** | 0.7 | 2.2 | **6.2\*\*** | 3.7 |
| **d) Growth** |  |  |  |  |  |  |  |
| % leaf area increase | **22.2\*\*** | **64.1\*\*** | **100.6\*\*** | 0.0 | **4.0\*\*** | 2.2 | 0.3 |
| % stem dia. increase | **22.2\*\*** | **75.7\*\*** | **29.6\*\*** | 0.0 | **6.6\*\*** | 2.4 | 1.9 |

**Figure 2:**
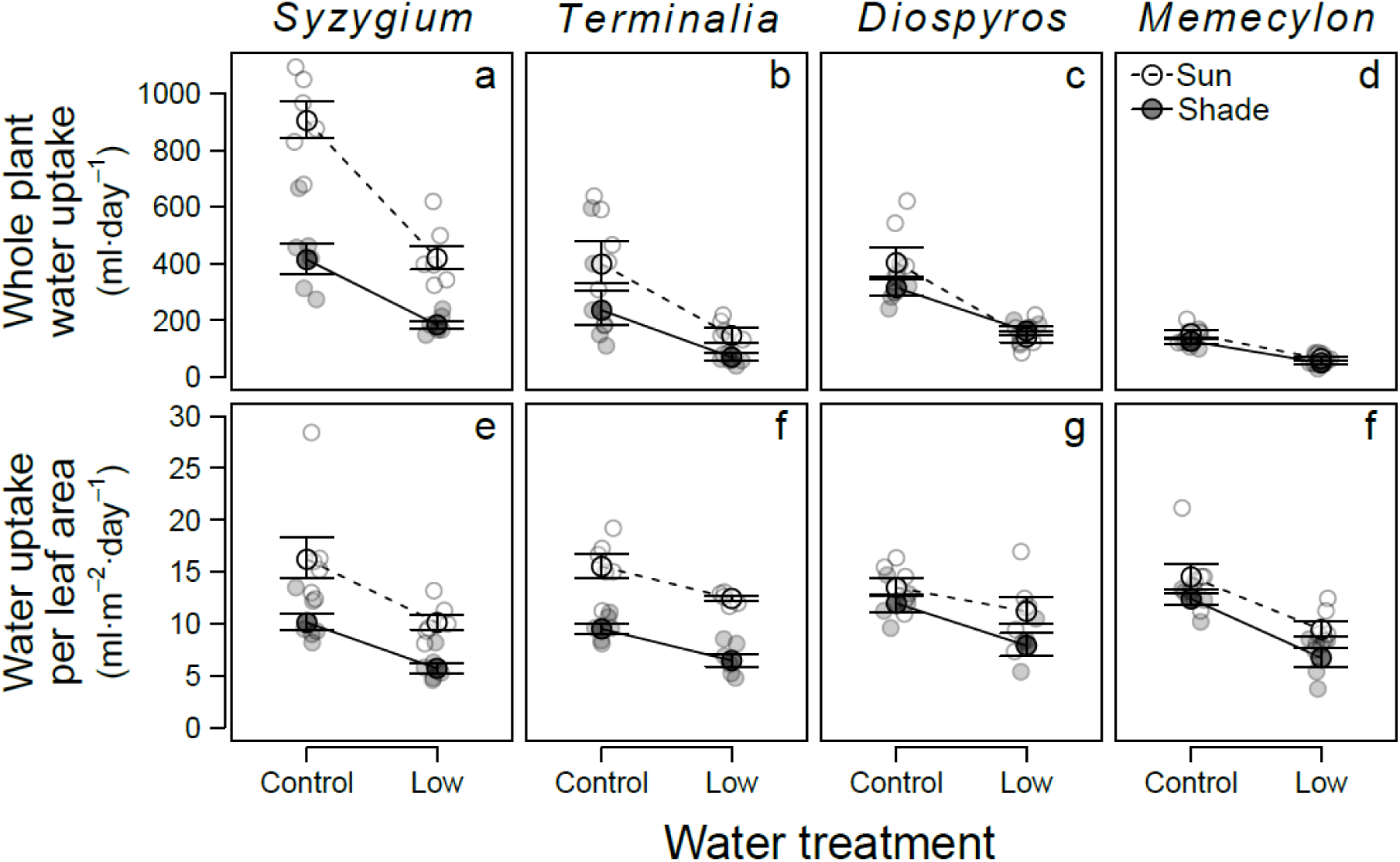
Water uptake for the study species in the experimental light and water treatments: a-d) average daily water uptake per plant; c-f) leaf area normalized daily water uptake. Error bars corresponds to ± 1 standard error. Open symbols and dashed lines represent plants in full sun light, and solid symbols and lines plants in the shade treatment.

Whole plant water uptake was qualitatively similar when water uptake was quantified on a leaf area basis (Fig. 2c-f, Table 2). There was no significant three way interactions, and the reduction in water uptake per leaf area between the open sun and shade treatments was lowest for Memecylon and highest for Syzygium. The decrease in water uptake per leaf area in the low water treatment was not dependent on light, and was similar for all four species as indicated by the lack of light into water, and species into water interactions, respectively (Table 2). Despite the overall congruency in whole plant water use and leaf area normalized water use, the decrease in the effect size in the water treatment implies a changes in leaf area allocation in response to the limiting water condition.

For the most part the affects of the light and water treatments on leaf physiology mirrored what was observed for plant water uptake (Fig. 3, Table 2). Photosynthesis, stomatal conductance and transpiration decreased in the shade and low water treatments, but we observed no interaction between the response to shade and water treatments (Table 2). As seen in water uptake, the change in photosynthesis and transpiration with shading differed among species. While the decrease in photosynthesis and transpiration was high in Terminalia, there was no significant change in photosynthesis for Memecylon in the shade.

**Figure 3:**
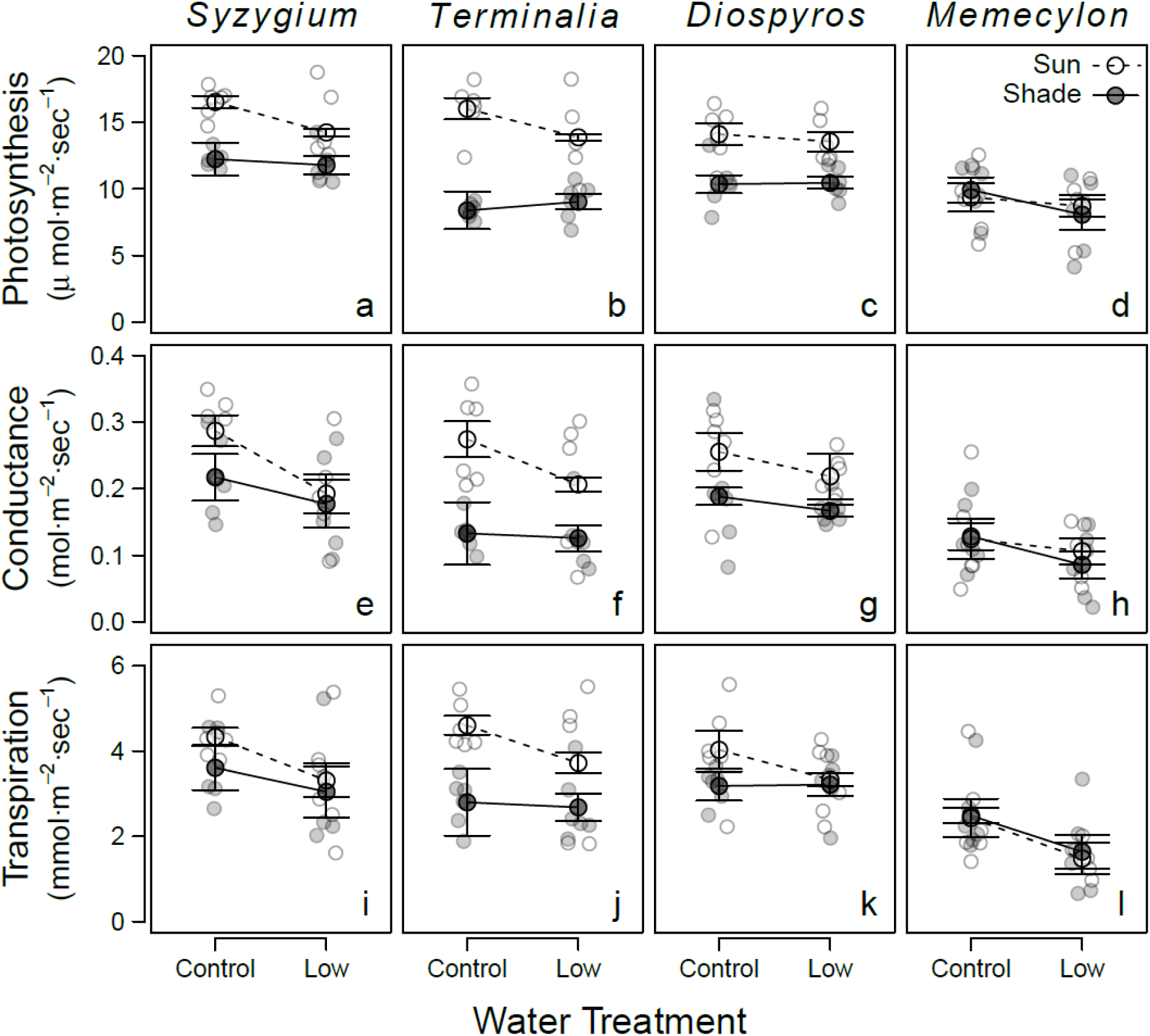
Response of leaf physiology in the study species to the light and water treatments: a-d) Photosynthesis (A_net_); e-h) Stomatal conductance (g_s_); and, i-l) Transpiration (E). Error bars corresponds to ± 1 standard error. Open symbols and dashed lines represent plants in full sun light, and solid symbols and lines plants in the shade treatment.

The results for leaf photosynthesis and stomatal conductance differed from what we observed for water uptake in an important way in that the magnitude of change was higher for the light relative to the water treatments. This is understandable in that both photosynthesis and conductance are expected to be strongly driven by light intensity. Additionally, unlike with water uptake we did not see a light into species interaction for conductance indicating that the effects of light were similar for all four species.

Light did not alter biomass allocation (Fig. 4, Table 2). The low water treatment resulted in a decrease in leaf mass fraction and a corresponding increase in root mass fraction in Diospyros and Terminalia, but not in Syzygium or Memecylon (Fig. 4, Table 2).

**Figure 4:**
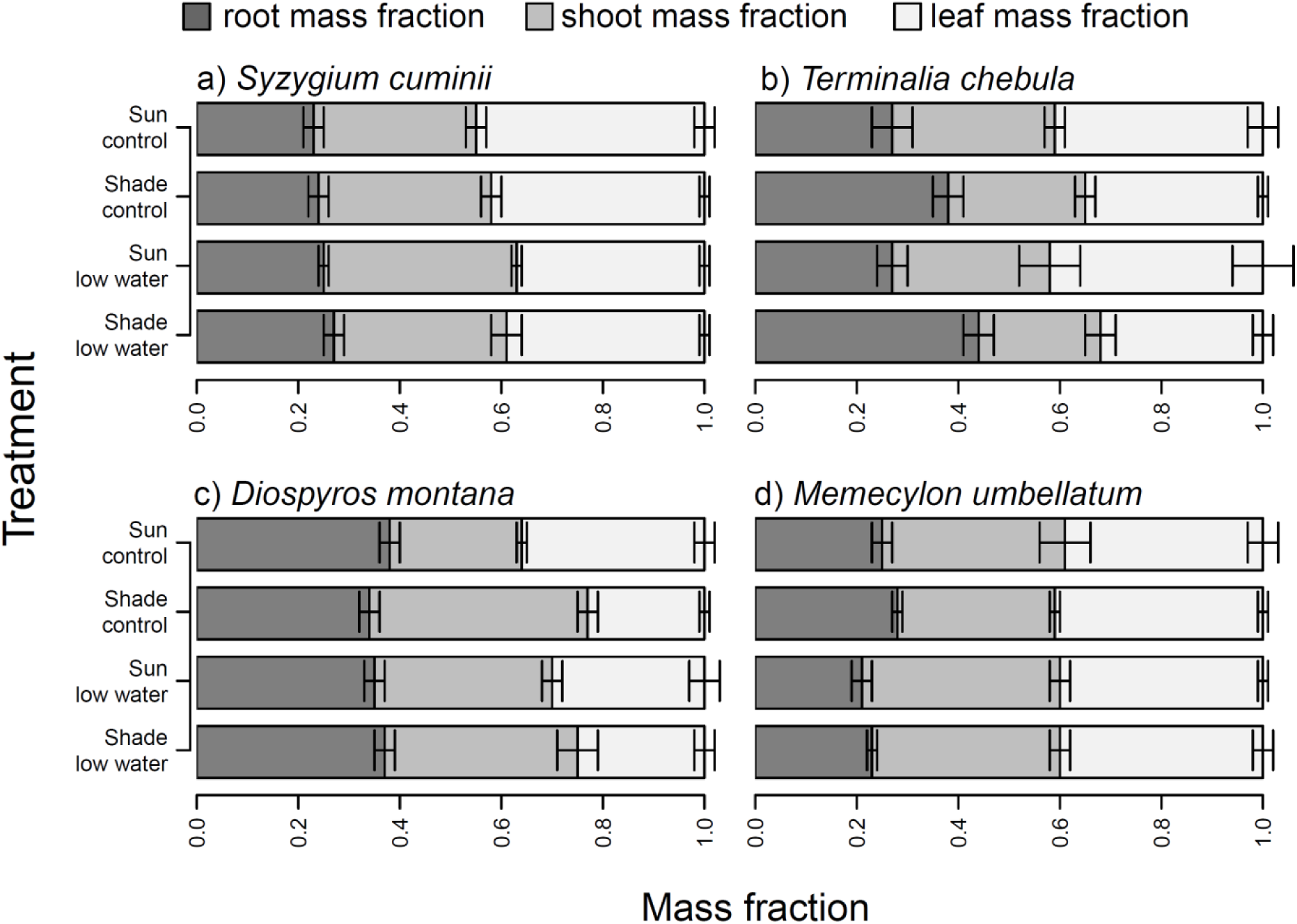
Biomass allocation to roots, stems, and leaves in response to the light and water treatments for: a) *Syzygium cumini*; b) *Terminalia chebula*; c) *Diospyros montana*; and, d) *Memycelon umbellatum*. Error bars corresponds to ± 1 standard error. Mass fractions for leaves, stems and roots are presented as the dry weight in these organs as a fraction of the total dry biomass. Root mass fractions are shown in dark gray; stem mass fractions in medium gray; and, leaf mass fractions in light gray.

Growth responses to the treatments in these four species were consistent with what was observed for plant water uptake and leaf physiology. The low water and shade treatments resulted in a reduction in the percent increase in leaf area and stem diameter (Fig. 5, Table 2). As seen with water uptake and physiology, the lack of a light into water interaction indicates that the shade treatment did not alleviate the reductions in growth caused by the low water treatment (Table 2). Mirroring what was observed in whole plant water uptake, species differed in how they responded to the shade but not the water treatments (Table 2). Terminlia and Memecylon showed the highest and lowest changes in growth, respectively.

**Figure 5:**
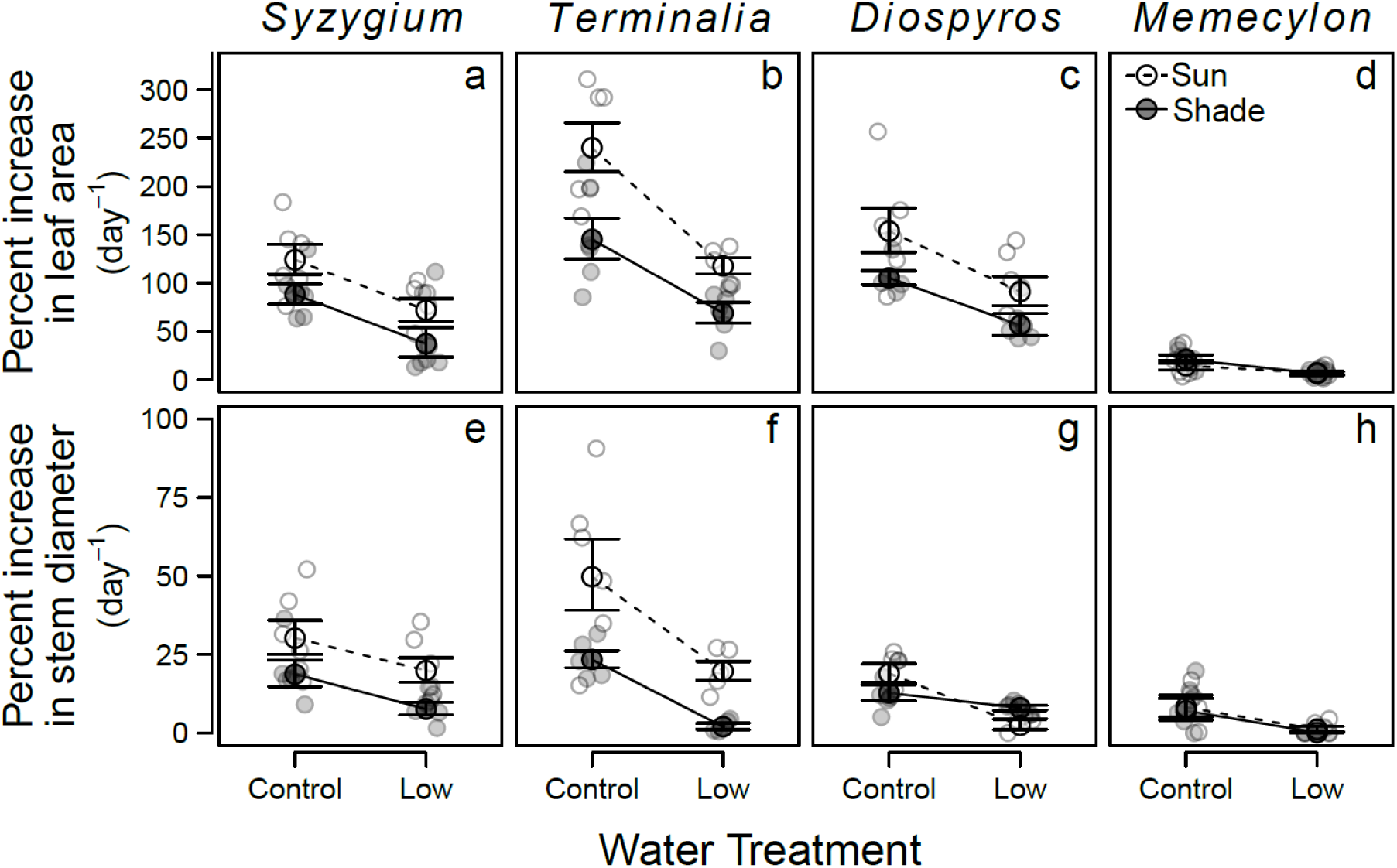
Growth measures for the four species in response to the light and water treatments: a-d) Percent increase in leaf area; and, e-h) Percent increase in stem diameter. Error bars corresponds to ± 1 standard error. Open symbols and dashed lines represent plants in open sun, and solid symbols and lines plants the shade treatment.

## Discussion

Whole plant water uptake varied dramatically in seedlings of these four co-existing species with a five-fold difference in maximum water uptake in well watered conditions in full sunlight. As expected species responded to low water availability by reducing water uptake, and this reduction was similar across species. Shading also resulted in reduced water uptake, but in a species-specific manner. The responses to the low water treatment were similar for plants in the sun and shade treatments, and this indicates that low light levels did not alleviate effects of reduced water availability. This also suggests the lack of a trade-off between responses to limiting light and water. Overall, responses of leaf physiology were congruent with what we observed for water uptake, but the relative effect of shade was larger for leaf physiology than water uptake. The reductions in water uptake observed in limiting light and water were reflected in decreased growth.

The large differences among species in whole plant water uptake in the full sunlight and well watered conditions was associated with a two-fold difference in photosynthesis, and 16-fold difference in growth as measured by percent increase in leaf area. However, despite these large inherent differences in resource use, we did not detect any differences in responses to limiting water treatments in these species. This was true for water uptake, leaf physiology and growth, and implies that the more conservative species did not fare better in limiting water conditions. This is contrary to suggested trade-offs in conservative strategies between performances in resource-abundant versus resource limiting conditions (Meinzer et al. 2010).

The lack of differences in how these species responded to limiting water conditions may be in part due to strong selective forces imposed by the highly seasonal climate and the long and severe dry season that species have to endure in this region. In contrast to the responses to limiting water, species responses to the shade treatment differed for water uptake, photosynthesis and growth. Memecylon which exhibited the lowest overall change in response to shade is the only species which is dominant in the closed forests, and the observed differences in shade tolerance are likely important in being able to establish in the edge and closed habitats with lower light availability.

The lack of a significant interaction between the light and water treatments for all of the study species (no significant three-way interaction between species, light and water) indicated that responses to limiting water and light were independent of each other. Thus, we did not find evidence for a negative relationship between responses to limiting light and water in support the proposed trade-off between these responses (Smith and Huston 1989, Holmgren et al. 2012, Kupers et al. 2019). Two of the study species increased allocation to roots and decreased allocation to leaves in response to the low water treatment. However, we did not observe increased allocation to leaves at the expense of allocation to roots in response to shade, which is expected to be one of the primary mechanisms proposed to explain the tradeoff between responses to limiting light and water (Smith and Huston 1989). The lack of an interaction between light and water also rules out positive relationships between responses to limiting light and water in these species, and suggests that shade did not ameliorate responses to drought (Holmgren 2000, Sack and Grubb 2002), and did not result in coordinated whole plant strategies (Reich 2014). Seedling responses to limiting light and water may be able to evolve independently of each other. This will allow wider ranges of morphological and physiological trait combinations in these species which would allow extensive niche differentiation to effectively explore the highly heterogeneous, seasonally and temporally variable light and water environments in these tropical dry forests.

How plant water uptake changed in response to limiting water and light treatments was largely congruent with change in leaf physiology as well as whole plant. However, there were important differences in the relative magnitude of effect sizes for the light and water treatments for the different responses examined. Not surprisingly, the effect of the water treatment was larger relative to the shade treatment for plant water uptake. The relative magnitude of the effect size for the water treatment decreased for leaf area normalized water use, and this indicates that species did respond to limited water availability by decreasing total leaf area. In contrast to plant water uptake, the relative effect of the light treatment was more pronounced for leaf level physiology. Importantly, the relative effect of the water treatment was more pronounced for growth, mirroring what we observed for plant water uptake and not leaf physiology. Thus, growth in seedlings of the study species, a measure of overall performance in the face of limiting water and light, was better reflected in whole plant water uptake than in leaf level physiology. The difficulty in quantifying whole plant level water use has often resulted in relying on organ level proxies (Dial 2004, Rodríguez-Gamir et al. 2016). This result highlights the importance of examining whole plant water use to understand plant responses to changing light and water conditions.

Several studies that have examined trade-offs in responses to limiting light and water have often used natural gradients instead of controlled experiments owing to the advantage of exploring patterns in a larger set of species (Kupers et al. 2019). While these studies are able to examine larger numbers of species, this comes at the cost of not being able to tease apart confounding correlated environmental factors that can influence species responses. While this problem is overcome in experimental studies like this one, the number of species examined in this study was limited, and thus, caution should be exercised in extrapolating these results to the larger community. The water and light levels selected for this experimental study were chosen to represent limiting but not lethal conditions to allow us to examine physiologically relevant changes in the responses of seedlings of these dry tropical forest species. These light and water conditions selected are ecologically relevant in that the limiting light levels are representative of levels in the understory environments in forests in this region, while the low water levels reflect water availability in these forests during the dry season. However, previous work has indicated the possibility of a non-linear responses to varying light and water conditions (Holmgren 2012). Thus, controlled experiments across wider gradients of light and water availability and more species are essential to disentangle the complex responses and associated trade-offs.

## Conclusion

We report large variation in whole plant water use in seedlings of four dominant coexisting species of a seasonally dry tropical forest. Our results demonstrate that seedling responses to limiting light and water conditions were independent of each other and this suggests that responses to limiting water may evolve independently of shade tolerance. This would allow larger potential combinations of light and water responses to allow these species to adapt to light and water niches in dry tropical forests. Growth responses in these species were better explained by whole plant water use than leaf physiology, and this highlights the importance of studying whole plant water use responses in co-occurring species to better understand species establishment as well as the complex processes of coexistence and community assembly in seasonally dry tropical forests.

## Contributions

RS, DB and AG designed the study. RS conducted the experiment with help from AJ, KMN, NMB, and AG. RS and DB performed the analysis, and wrote the manuscript with input from AJ, KMN, NMB, and AG.

## Acknowledgements

We thank Nitish Lahigude for his help in propagation and maintenance of plants; Vivek Broome of J.E. Farms, Pune for providing saplings.

## Notes

### Competing Interest Statement

The authors have declared no competing interest.

