## Supplementary Data for "Response of whole plant water use to limiting light and water conditions are independent of each other in seedlings of seasonally dry tropical forests"

**Supplementary figures**

*List of Figures*

Figure S1: Map showing location of the garden experiment in Pune, and the field site.

Figure S2: Climate data for the region.

Figure S3: Relative light intensity across the three habitat types in Nigdale, Maharashtra.

Figure S4: Gravimetric soil moisture content in the habitat types Nigdale, Maharashtra.

Figure S5: Pot water content for control and low water treatments and light levels for the shade treatment and sun treatments.

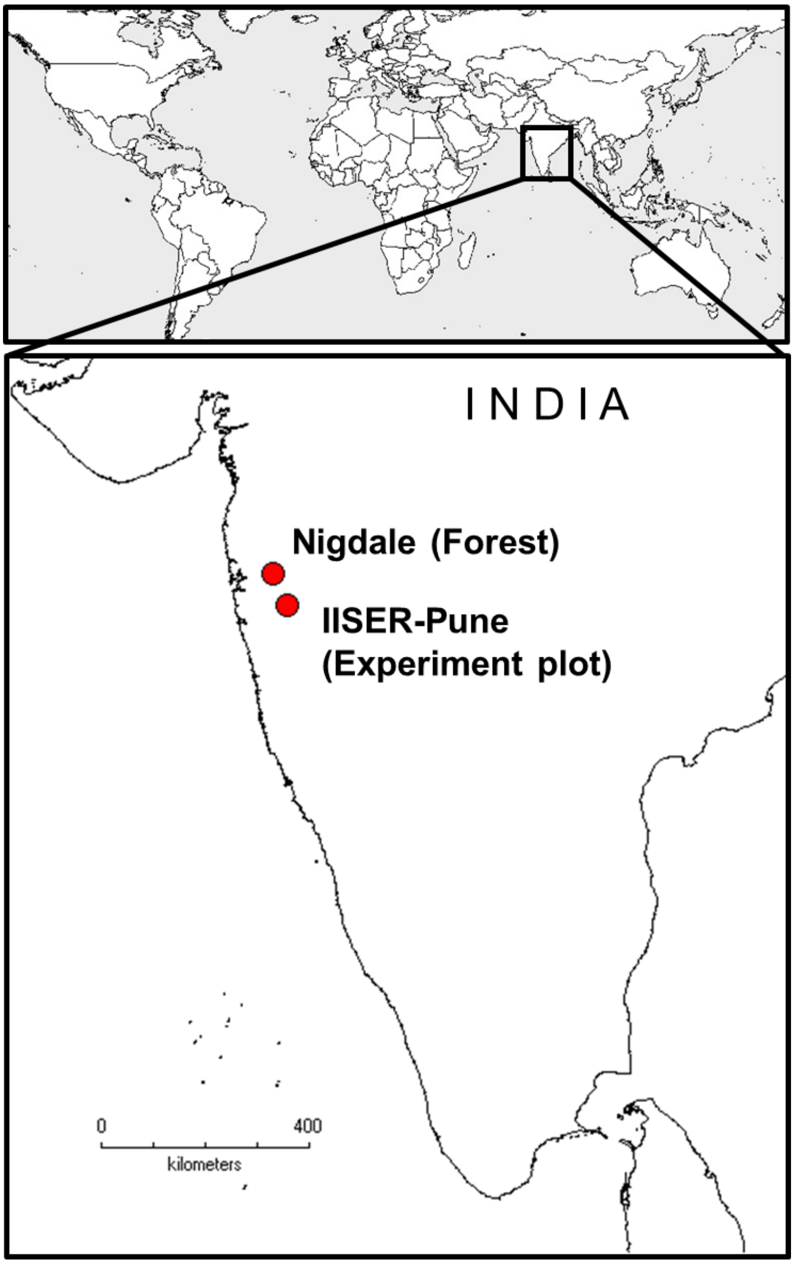


**Figure S1:** Map showing location of the garden experiment in Pune, and the field site from where species distribution data was collected in Nigdale, Maharashtra located at the northern end of the northern Western Ghats.

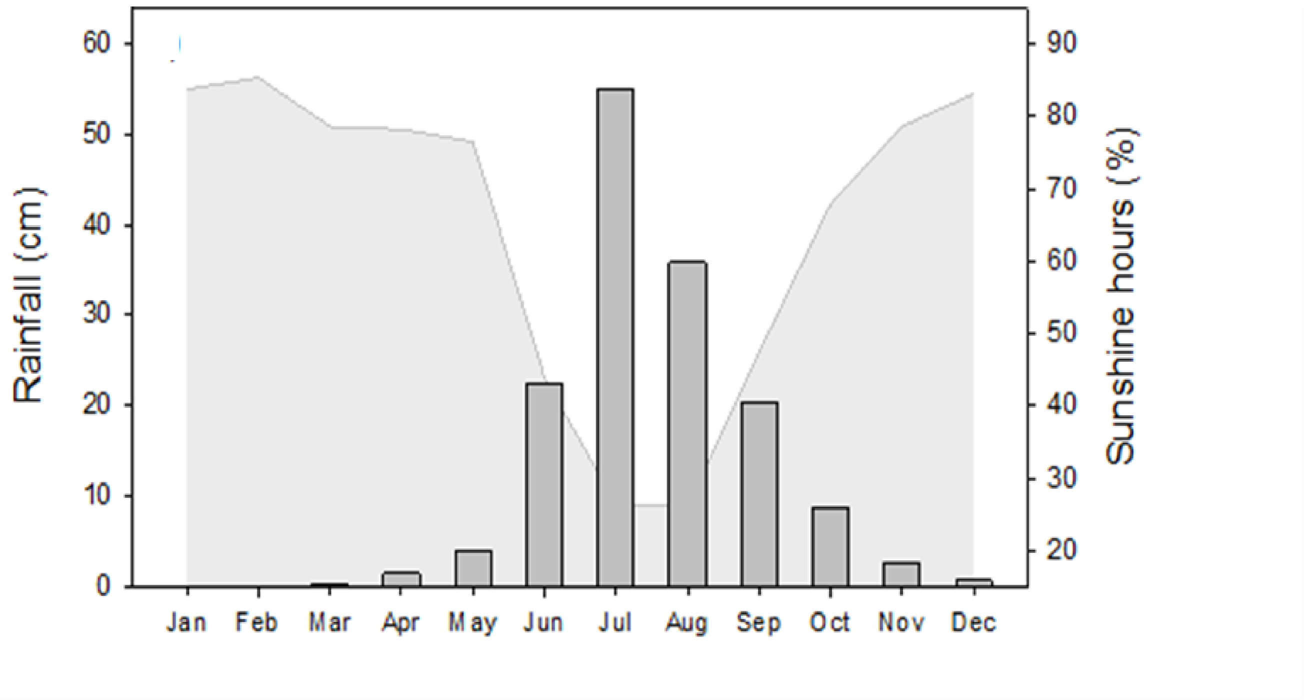


**Figure S2:** Climate data for the region from where species are abundant (Pune, Maharashtra, India). Monthly averaged precipitation (1961-1990) - Dark grey vertical bars; and, sunshine duration (light grey curve) are from a high resolution global dataset (New et al. 2002).

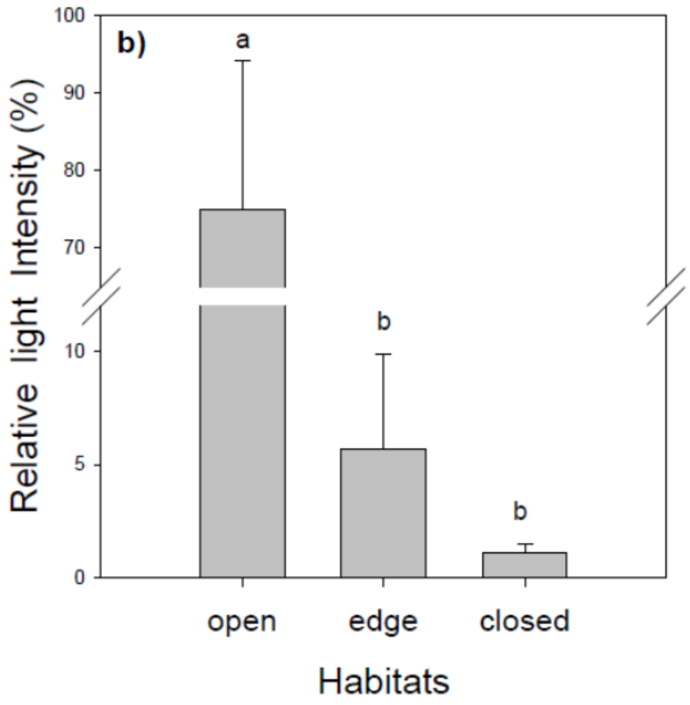


**Figure S3:** Relative light intensity across the three habitat types (Open fragmented forest patches-grass matrix; transition forests - Edge; and, Closed canopy forests) in Nigdale, Maharashtra in the Northern Western Ghats. This information was used to determine the intensity of the shade treatment.

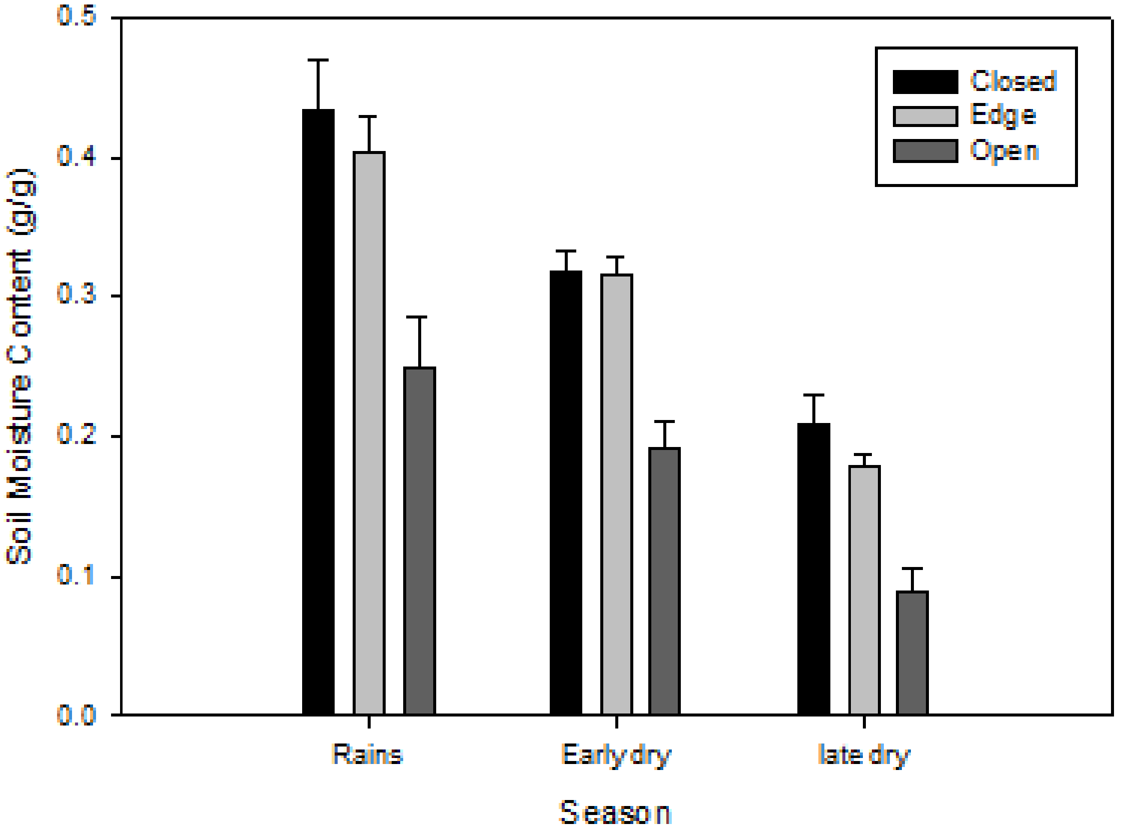


**Figure S4:** Gravimetric soil moisture content in the habitat types (Closed canopy forests; transition forests - Edge; Open fragmented forest patches-grass matrix) in Nigdale, Maharashtra in the Northern Western Ghats. Measurements were made during the latter part of the rainy season (September); early dry season (December); and late dry season (April) of 2015. This information was used to decide the watering treatment.

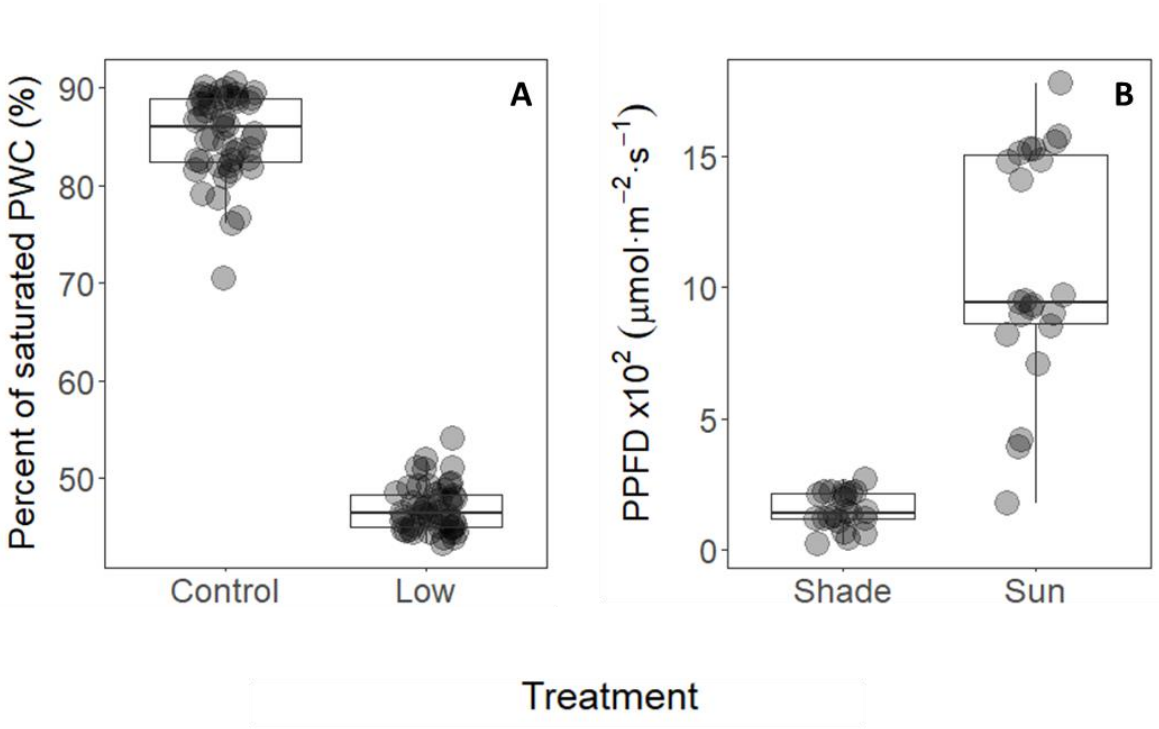


**Figure S5:** Treatment conditions imposed during the 50 day experiment period. A) Water treatment in terms of pot water content (PWC) corresponding to control (mean= 85% of saturated PWC, mode = 95%) and limiting water level (mean=46%, mode= 43%). Each point corresponds to the PWC of a single pot averaged over the treatment period. The control pots were reset to 95% while the low water treatment pots were reset to 45% every 3rd day. B) Photosynthetically active radiation measures for the shade treatment and sun treatment conditions in terms of average photosynthetic photon flux density (PPFD) measured during the course of the experiment. Each point corresponds to the PPFD of a day measured at 1230 Hrs.
